# Operationalising the context in regenerative agriculture: decision-making and farm variability

**DOI:** 10.64898/2026.09.28.748194

**Authors:** Tuomas J. Mattila, Jari Niemi, Henri Kajasilta, Tanja Myllyviita, Tommi Vippola, Layla Höckerstedt, Sampo Pihlainen, Johan Ekroos

## Abstract

Soil degradation is a widespread challenge that requires a broad response at the individual farm level. To ensure effectivity, the practices should be tailored to the farm context: land manager objectives and farm specific challenges. These have however been difficult to quantify. Here we demonstrate that a workable farm context can be created based on a value survey, open satellite and soil data, and published models for vegetation gross primary productivity and soil erosion. Based on the findings, despite individual differences, farmers value profitability and operational efficiency, but also biodiversity and soil health. At least the regenerative farmers surveyed also value working for the greater good more than maintaining tradition or power. In spite of wide differences in farm production orientation, we also found that each farm also had a broad variation in individual fields GPP. Most fields have a stable GPP level, which is either high or low, and that there is a 2–3-fold difference between the weakest and best producing fields indicating the potential for improving GPP by improving the growing conditions on currently weak fields. In addition, soil loss was found to be highly concentrated in critical source areas, where 10% of the field area contributed to 50% of the soil loss. Overall, open data can be linked to modelling workflows to rapidly produce a decision-making context for farmers. This facilitates benchmarking and co-learning as well as enables land managers and advisors to identify the farm context for planning effective responses to soil degradation.

## 1. INTRODUCTION

Soil degradation impacts over 52% of European farmland (Prăvălie et al., 2026) and is forecasted to worsen with climate change (Afshar et al., 2026). Farmers have responded to this challenge by adopting regenerative agriculture, a farm management style where production goals are increasingly balanced with soil and ecosystem improvement goals (Beacham et al., 2023; Jaworski et al., 2024; Schreefel et al., 2020). These goals are pursued by minimising tillage, increasing vegetation cover, and increasing crop rotation diversity (Emmett et al., 2025; General mills, 2019; Williams, 2022). The practices can result in multiple benefits across objectives (Liu et al., 2019; Mattila, 2024; Qiao et al., 2022; Winberg et al., 2026). However, they can also reduce production or operational efficiency (Emmett et al., 2025; Pannell et al., 2006; Schreefel et al., 2022). How to balance trade-offs between the multiple objectives, when they cannot be improved simultaneously?

Applying regenerative agriculture requires tailoring it to the farm context, the broad decision-making environment, which consists of both farm properties and farmer priorities (General mills, 2019; Williams, 2022). Values and priorities influence the acceptability of management outcomes and trade-offs (Beacham et al., 2023; Hill and Bradley, 2024; Jaworski et al., 2024). Farmers already balance complex orientations, ranging from purely instrumental to the intrinsic value of work, expression through creativity, and social networks building (Hill and Bradley, 2024). The relative importance of these orientations is linked to the decision context and to the intrinsic values of the farmers (Pannell et al., 2006). Farmers have traditionally been considered conservative and profit-oriented, but recent large-scale surveys have challenged this notion, instead portraying a diverse range of farmers who value novelty and self-direction as well as universalism and benevolence (Beacham et al., 2023; Graskemper et al., 2022; Sorvali et al., 2022). With the great agricultural shift in the workforce, farming has transitioned from everybody’s occupation to the work for the common good (Graskemper et al., 2022), which is well aligned with the community, soil, and ecosystem restoration goals of regenerative agriculture (Beacham et al., 2023).

Several indicators have been developed to track soil functions and farm performance over time (Donovan et al., 2025; Panagos et al., 2025; Qiao et al., 2022), but the indicators are fairly complex. For example, a recent consensus indicator list for tracking farm performance comprises of 24 indicator across eight attributes, such as producer well-being, resource reserves, and productivity (Donovan et al., 2025). Obtaining data for all the indicators is difficult, especially if they should be scaled across farms containing tens of fields and hundreds of field management units. Focusing on only a few indicator fields in each farm risks missing fields or field parts, that have localised soil degradation processes (Mattila and Vihanto, 2024). For erosion and sediment loss, it is well known that 5-20% of area typically contributes to 50-80% of the soil loss (Mattila and Niemi, 2026; McDowell et al., 2025). To identify and focus on the critical source areas, these would have to be identified at the farm level using high-resolution monitoring.

High-resolution data enable a more effective use of financial compensation to support transition (Ferraro, 2008; Latacz-Lohmann and Van der Hamsvoort, 1998). Many methods for determining compensation levels depend on the land managers having high-quality information on their field productivity and improvement potential (Boufous et al., 2023; Ferraro, 2008). Although remote sensing can provide such information, it needs to be translated into interpretable outcomes through modelling. For example, models have been developed for vegetation primary productivity (GPP) (Vira et al., 2025), soil loss (Räsänen et al., 2024), and pollinator abundance (Abdi et al., 2021; Blasi et al., 2021; Gardner et al., 2020). However, this work is still ongoing for most of farm performance indicators.

It is challenging to balance multiple indicators with variable farmer preferences and on-farm situations. With more indicators, the probability of conflicting outcomes increases. This accentuates the need to effectively resolve trade-offs to avoid analysis paralysis. Regenerative agriculture tries to work around this by recommending context-specific solutions (General mills, 2019; Williams, 2022), where practices are tailored to the farm situation and the land managers’ goals and values. However, this leaves open a set of key questions: How do we define the context? What are the critical differences between farms and how should these differences be determined? What farms share similar goals and soil quality challenges and can therefore learn from each other?

In this study, we demonstrate how on-farm contexts can be operationalized through a simple values questionnaire combined with indicators derived from open data. We explored farmer priorities and values to investigate how they balance profitability and operational efficiency with long term soil health and biodiversity goals. To benchmark farms and to identify potential for improvement, we classified fields based on their satellite-estimated long-term productivity and stability. Finally, we mapped the erosion and sediment loss at the sub-field level to identify critical areas requiring urgent soil conservation practices to avoid soil degradation. The farmer values, field productivity, and localized soil loss maps present a first estimate for a generalisable farm context. Overall, the decision-making criteria prioritized by farmers could be linked to existing methods to monitor changes in farm performance over time. Our findings demonstrate that the vague concept of an on-farm context can be operationalised into an actionable set of farm-relevant indicators.

## 2. MATERIALS AND METHODS

### 2.1 Description of the participating farms

We tested the approach on 10 volunteer farms, which were divided into dairy farms (n=4) and grain farms (n=6). The farms were in an area of 220 × 390 km from Kymenlaakso to North-Ostrobothnia, spanning the main cropping areas of Finland. Typically to Finnish croplands the farms soil types were dominated by Cambisols (71%), Regosols (10%), Podzols (7%) and Leptosols (7%), but with only 1% of Histosols. The participating farms were two to ten times larger than the Finnish average farms, averaging 270 ha of field area (122-560 ha range). Their fields (n=852) were also twice the size (3.9 ha) of the average fields. Both farm and field size represented the upper quartile of production for grain farms and dairy farms, indicating that the farms were highly productive professional farms (Natural Resources Institute Finland, 2026). Three of the grain farms and one dairy farm were in organic production, representing a much larger share (40%) than the 13% Finnish average (Finnish Food Authority, 2026). The crop rotations were diverse, with annual/perennial crop rotations in 60% of the area, and cereal or grass monocropping in only 11% of the field area. This diversity was driven by the high share of organic farming, which had 92% of land area under perennial/annual rotation. The conventional farms had 39% in perennial/annual rotation, 35% in diverse annual rotations, and 15% under monocropping; the share of diverse rotations was also much larger than the Finnish average of 10% (Peltonen-Sainio and Jauhiainen, 2019).

The dairy farms had 2-3 managers each, whereas the grain farmers operated the farms by themselves. The average age of the farmers was 48 years (range 29-65; Finnish average 55 years), and they had an average of 17 years of experience in managing their farm (range 0-40 years). All farmers had relevant official education (agriculture or engineering) for the occupation (13% university, 47% applied sciences, and 40% vocational upper secondary). Overall, the participating farmers were skilled, well educated, and motivated farmers, who operated and actively developed their relatively large farms.

### 2.2 Assessing values and decision-making criteria

We evaluated farmer values and decision-making criteria through two widely used methods: the Schwarz value questionnaire used by the European Social Survey (ESS) and the simple multi-attribute rating technique (SMART) (Winterfeldt and Edwards, 1993). The ESS questionnaire maps respondents’ values to a universal framework of 10 basic values: universalism, benevolence, security, power, hedonism, achievement, stimulation, self-direction, tradition, and conformity. These values were mapped using the most recent questions of the 11^th^ round of ESS (ESS, 2024). The survey questions enabled comparison to country-wide responses and earlier farmer-oriented surveys (Graskemper et al., 2022; Sorvali et al., 2022).

The SMART weight assessment focused on detailed, actionable decision-making criteria using a two-step hierarchical weighting system. The first step involved five broad scales of objectives for farm performance: social recognition, profitability, operational efficiency, future proofing and next generations, and a diverse and aesthetic environment. Each objective was split into four quantifiable subcriteria, resulting in a value tree with 5 objectives and 20 subcriteria. The respondents were asked to first rank the top-level objectives against each other by giving the most important objective 100 points and then giving the other objectives lower scores, which relate their importance to the most important objective (i.e. SMART weighting). The respondents then went through the subcriteria with the same ranking process. Finally, they reassessed the top-level objective weights after examining the subcriteria’s calculated overall weights. In addition, the respondents were asked to state their familiarity with the subcriteria on a Likert scale of 1-4 (1: unknown, 4: in routine use). This enabled highlighting high-weight and low-familiarity subcriteria for further research to improve their usability.

### 2.3 Classifying variability in the field productivity

To identify low- and high producing fields, we classified them based on gross primary productivity (GPP) and stability of GPP over time. GPP was estimated using a new model from satellite observations of vegetation cover combined with incoming solar radiation (Vira et al., 2025). In brief, we obtained 10 × 10 m multispectral imagery from Sentinel-2 (ESA, 2023), filtered the data for cloudiness, calculated a vegetation index (chlorophyll index, red-edge, CI_RE_=B7/B5-1) and multiplied this with daily photosynthetically active radiation (PAR), obtained from Copernicus Atmospheric Monitoring Service (CAMS) (Copernicus, 2026). This multiplication result was then used to estimate the daily GPP using Vira et al.’s (2025) statistical regression model. The 10×10 m resolution daily GPP was aggregated to the level of individual fields using the public field boundary data from the Finnish Food Authority (2026) and then aggregated to annual GPP for each field for each of the years 2020-2024.

The annual GPP was used to classify the fields based on productivity (higher or lower than average) and stability (coefficient of variation between years). We used the common 20% cutoff limit for high variability (Pimentel-Gomes, 2008). We also checked the data for trends using a least squares regression against years for each field with the R programming environment (R Core Team, 2024). Trends were recorded only for fields with statistically significant slopes (p<0.05). To explain the differences in GPP, the fields were classified by farm and production system, and compared with maps of depth-to-water (Salmivaara, 2020). The depth-to-water for 1 ha runoff accumulation limit (Salmivaara, 2020) was averaged for each field using QGIS (Dawson et al., 2026) and correlated with the GPP in R (R Core Team, 2024).

### 2.4 Prioritizing critical source areas for erosion and sediment loss

To study how erosion potential and sediment loss are distributed on each farm, we constructed 10 × 10 m erosion and sediment loss maps. The starting point for the mapping was a publicly available erosion susceptibility map (Räsänen, 2021), which was based on the RUSLE erosion model (Panagos et al., 2015), but excluded the cropping (C) and tillage (P) factors. We included these factors using field crop data for 2020-2025 (Finnish Food Authority, 2026), crop-specific, locally calibrated crop factors (Räsänen, 2024), and an Sentinel-2 (ESA, 2023) based estimate of soil cover and vegetation. Soil cover was ranked into three classes, using the Bare Soil Index (BSI), and corresponding RUSLE P-factors were assigned to each class: low soil cover, BSI > 0.1, full tillage P 1.0; medium soil cover, 0.021 < BSI < 0.1, minimum tillage P 0.75; and high soil cover, BSI < 0.021, no-tillage P 0.44. These P-factors were then adjusted with the living vegetation cover: if the NDVI was higher than 0.44, the P factor was reduced by 20%, in accordance with the average erosion reduction benefit from cover crops (Panagos et al., 2015) or vegetation in general (Räsänen, 2024). The class boundaries for BSI and NDVI were based on previous studies (Castaldi et al., 2023; Diek et al., 2017). However, we also tested the boundaries on fields with known tillage histories and time series of tillage events to ensure that they also applied to local conditions. We focused on the first three weeks of October, which is the end of the growing season and a time when tillage is mostly complete. To account for late tillage, we used the upper quartile BSI for evaluating soil cover and the average NDVI for vegetation in determining erosion reduction. The total erosion estimate was then calculated by multiplying the RUSLE erosion susceptibility with the yearly cropping and land cover specific C and P factors.

The erosion estimate did not describe whether the eroded material would result in soil loss at the field level. To estimate this, we calculated a sediment transport ratio (SDR) using a LIDAR-derived surface flow model (Forest Centre, 2024) to obtain the slope, catchment area, and distance to waterway for all field pixels (10 × 10 m resolution). These were then converted to an SDR using the equations of Räsänen et al. (2024). For analysis, we calculated the cumulative sediment loss curves and identified the fraction of fields responsible for 50% and 80% of the total sediment loss.

## 3. RESULTS

To investigate the critical differences in farm contexts, we first evaluated farmer values, objectives, and decision criteria weights. Then, we assessed trends in field productivity and consistent productivity differences. Then, we compared productivity with erosion and sediment loss. Finally, we evaluated the decision-making criteria, how they are used, and which criteria are good candidates for further operationalisation.

### 3.1. Values and decision-making criteria

To quantify farmers’ values and decision-making criteria, we asked them to evaluate the use and relative importance of different criteria. Gross profit was uniformly weighted as the most important criterion (Fig 1). The relative importance ranged from 7% to 29% of the total weight (average 15%), depending on the farmer. Similarly, almost all farmers ranked maintaining history and tradition as one of the least important criteria (0-3%). Other criteria weights, such as capital productivity, avoiding work peaks, peer support from other farmers, and the importance of food production, were more widely distributed. Apart from profit (15%), clear process organisation (8%), farming conditions for the next generation (7%) and soil health (6%) had moderately high weights. Overall, the farmers valued multiple criteria but prioritised profitability and operational efficiency.

**Figure 1.**
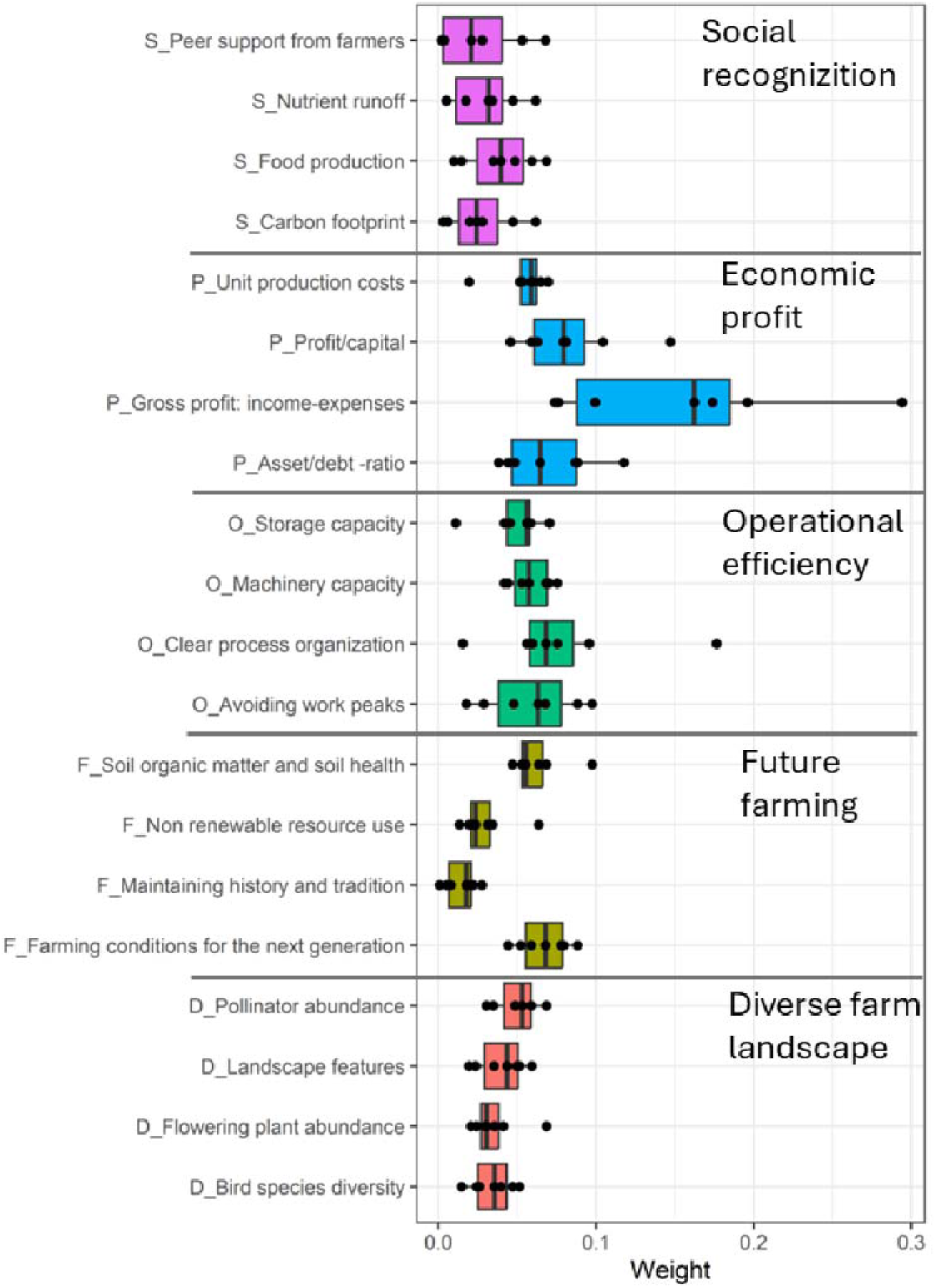
Decision making criteria and their weights for the five broad objectives. The sum of the weights of each criterion is set to 1 for each respondent.

The prioritisation of decision-making criteria correlated with the respondents’ values. Power was strongly correlated with gross profit (r=0.68), profit/capital (r=0.63), and unit production costs (r=0.60). Universalism was correlated with pollinators (r=0.62), soil health (r=0.58), and landscape features (r=0.55). Overall, the farmers showed a high preference for openness and self-transcendence and low preference for power, tradition, or achievement (Fig S1).

### 3.2. Variability in field productivity

To evaluate the productivity differences, we modelled gross primary productivity (GPP) for each field for 5 years and evaluated GPP stability and trends. Based on the findings, 65% of all fields had stable GPP, but very different average productivity (Fig 2). Approximately 37% of the fields were classified as high productivity, 28% low productivity, and 35% variable productivity. There was a weak, but significant correlation (R^2^=0.06***) between interannual variability (CV) and average GPP, suggesting that more productive fields are more stable. The largest variability was caused by two poor cropping years in 2021 and 2022, which had 8-10% lower GPP across fields. This reduction was concentrated on conventional grain farms (n=3), while the organic grain farms had no reduction in GPP. Dairy farms (n=4) had only 5% GPP reduction in the poor cropping years. Very few of the fields had a consistent trend in productivity (3% had a negative and 9% a positive trend). The stable productivity level was not explained by field water availability (i.e. depth-to-water), but more by farm management. Dairy farms had 16% higher GPP than grain farms and a diverse crop rotation was associated with higher productivity and lower variability (Fig S2). Nevertheless, each farm had a mix of low and high productivity fields. This suggests that improving poor fields and co-learning from best performing farms are valid strategies for improving farm productivity.

**Figure 2.**
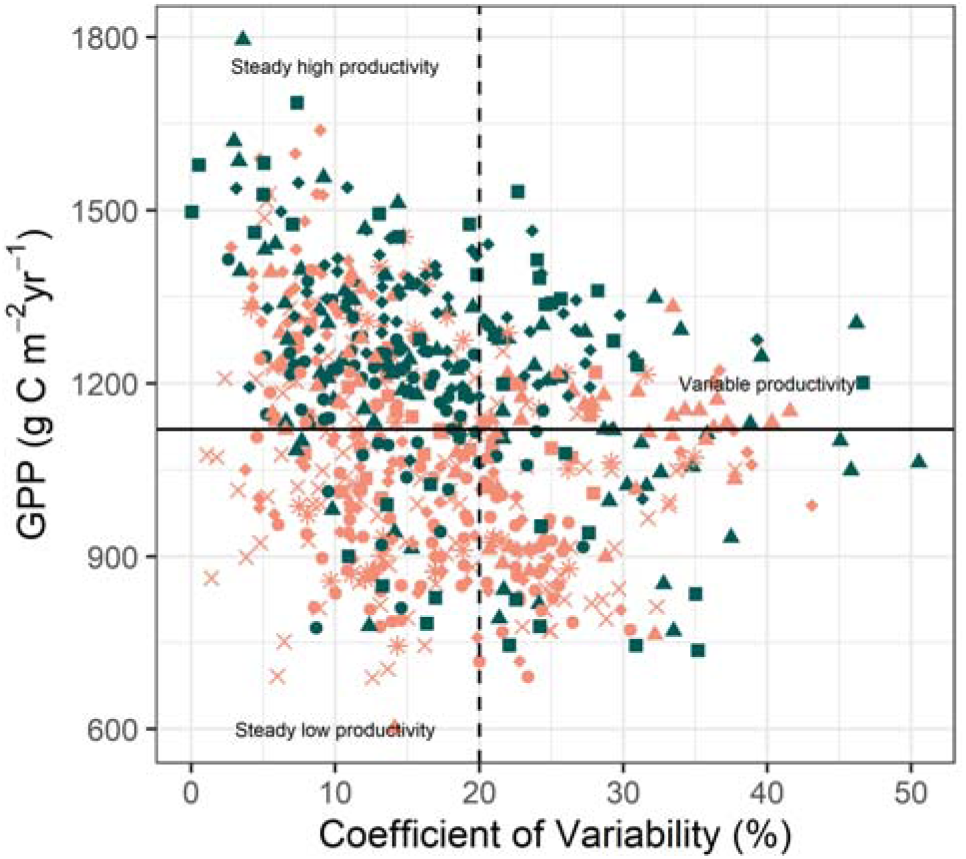
Average gross primary productivity (GPP) and interannual variability over five years in the studied dairy (green) and grain (red) farms. Symbols represent fields (n=852) of individual farms (n=10).

### 3.3. Critical areas for erosion and soil loss

To prioritize areas for soil conservation, we estimated erosion from soil susceptibility, crop choices, and tillage and combined these into high resolution erosion maps (Fig 4). Overall, the estimated average erosion was low (340 kg/ha/yr, Fig 3) and well below the soil formation rate (< 1 t/ha/yr). However, 5% of the area exceeded the soil formation rate, and 0.1% had a critical erosion rate (> 5 t/ha/yr). The generally low erosion rate was consistent with the farmers already having cropping and tillage practices, which had reduced erosion by 55% when compared to a situation with annual cropping and bare soil. These practices were not sufficient for soil conservation on 5% of the area, which should be targeted for further actions.

**Figure 3.**
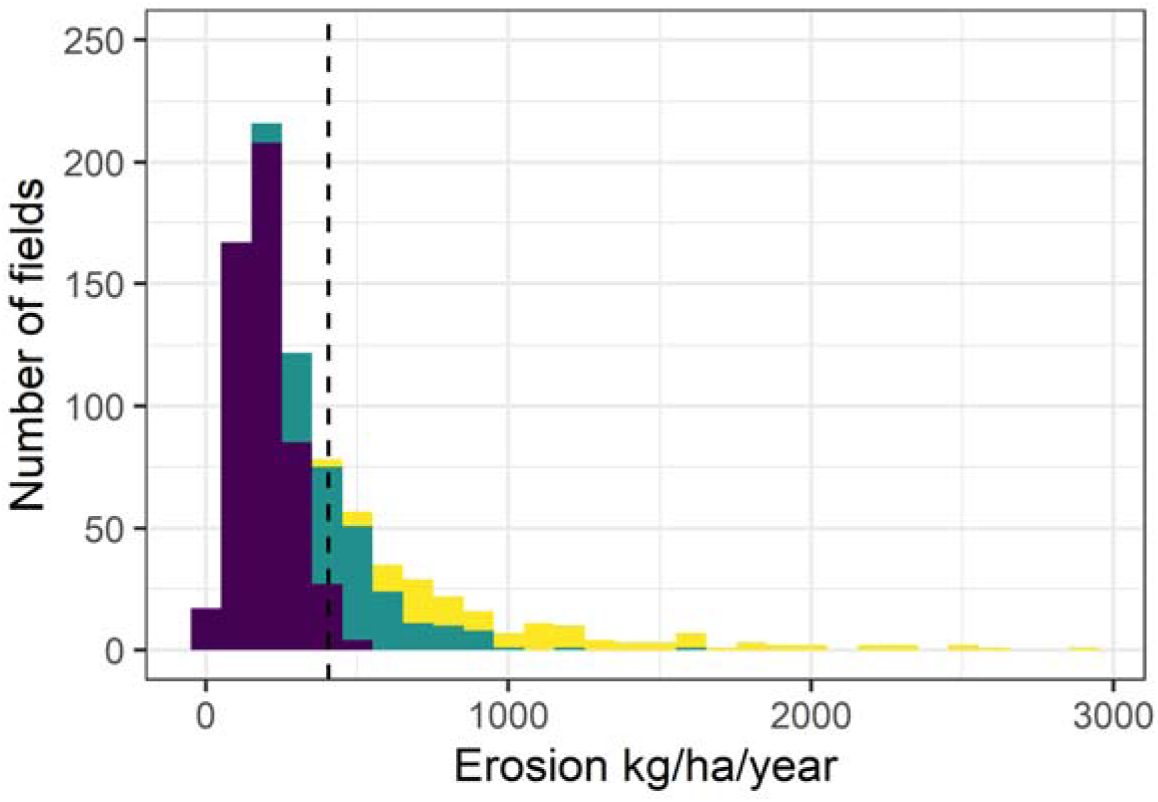
The distribution of erosion across the studied fields. Colors = yellow 50%; green 30%; and blue 20% of cumulative soil loss.

**Figure 4.**
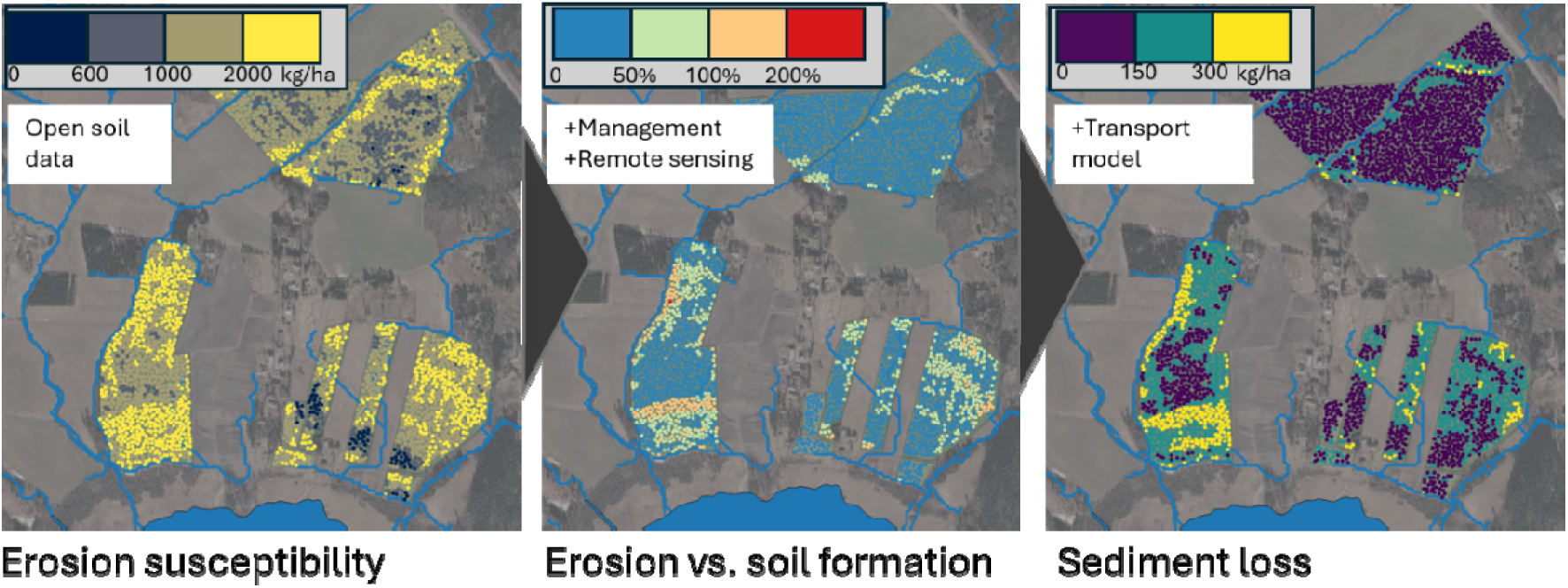
Example of the spatial distribution of erosion risk and sediment transport in one of agricultural catchments under study.

Erosion does not always result in soil loss from the field if the eroded material is deposited on field. To estimate the actual soil loss, we considered the proximity to concentrated flow paths and applied sediment transport ratios (SDR). We found that 50% of the soil loss happens over 10% of the field area, and that 80% of the loss happens on 33% of the field area (Fig 3). These critical areas are a priority for both soil conservation and waterbody protection from sediment load. On average, 31% of the eroded material left the fields through runoff and became sediment load. The average sediment load per farm was variable (38-266 kg/ha/yr) and strongly correlated with erosion susceptibility (i.e. slope position and soil texture) (R^2^=0.72***). Overall, erosion was a critical requirement for subsequent transport. Neither erosion nor soil loss correlated with GPP, indicating a low productivity but high erosion subgroup of fields that would be an ideal target for improved soil management. These target fields were characterized by annual cropping and low soil cover. Prioritizing these areas could simultaneously improve productivity and stop further soil degradation.

### 3.4. Workflows for operationalizing decision-making criteria

All of the farmer decision making criteria in Fig 1 could not be operationalized in this study. However, considered the availability of data and models for each criterion (Table 1). Each criterion was also evaluated by its importance and familiarity to identify targets for further model development. We found that although models and data are available, the amount and accessibility of data differ, as does the ease of use of the methods. Some methods, such as biodiversity or preservation of history, require on-site monitoring and extensive surveys, while others can be done with remote sensing products. Most of the responses indicated a polarised use of criteria: they were either “not used at all” or “used routinely”. For example, financial accounting methods were important, routinely used by most participating farmers, and well accessible. However, operational efficiency criteria were highly important, but much less utilised, perhaps because their interpretation would need access to multiple farms’ data for benchmarking.

**Table 1.**
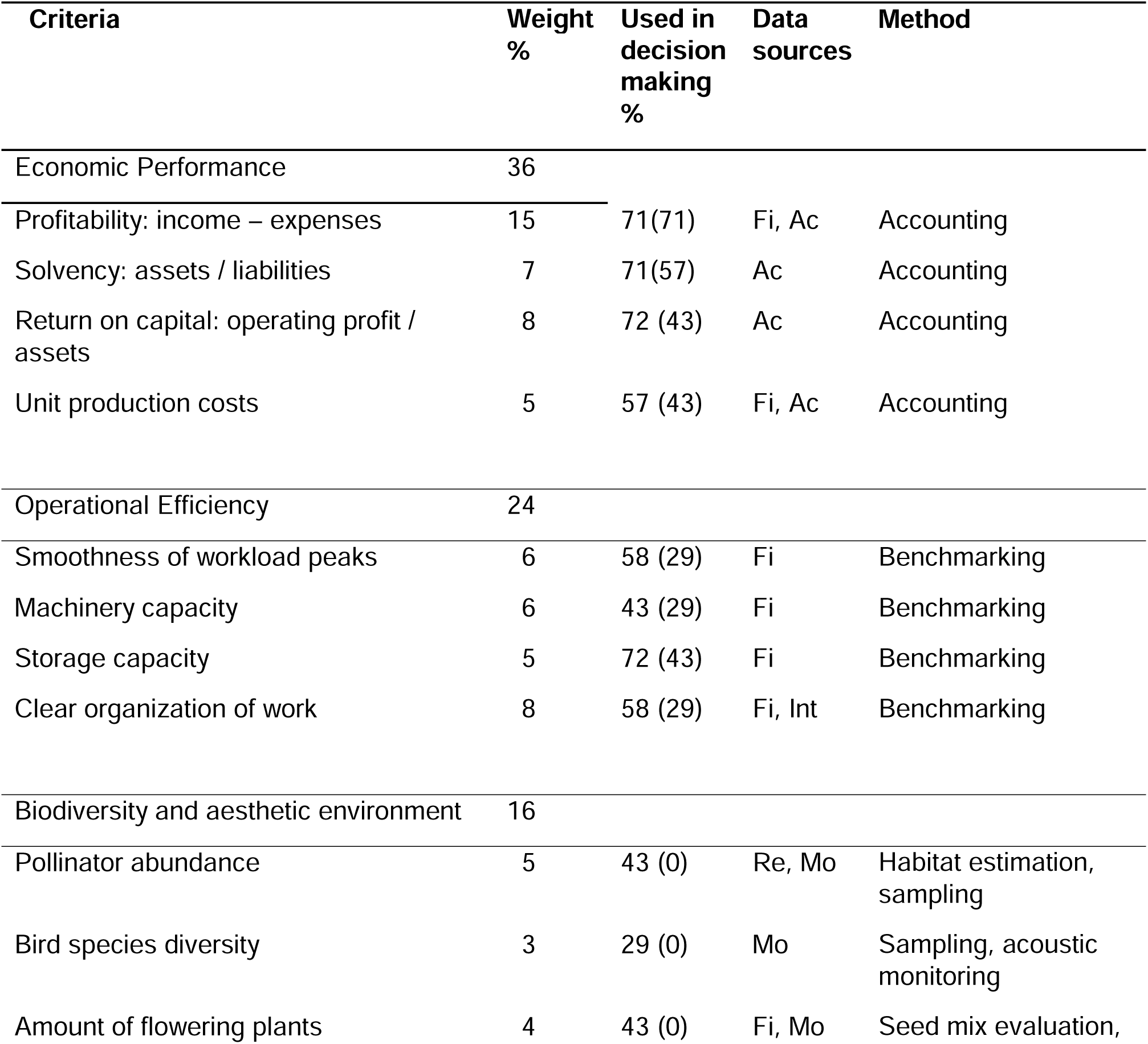
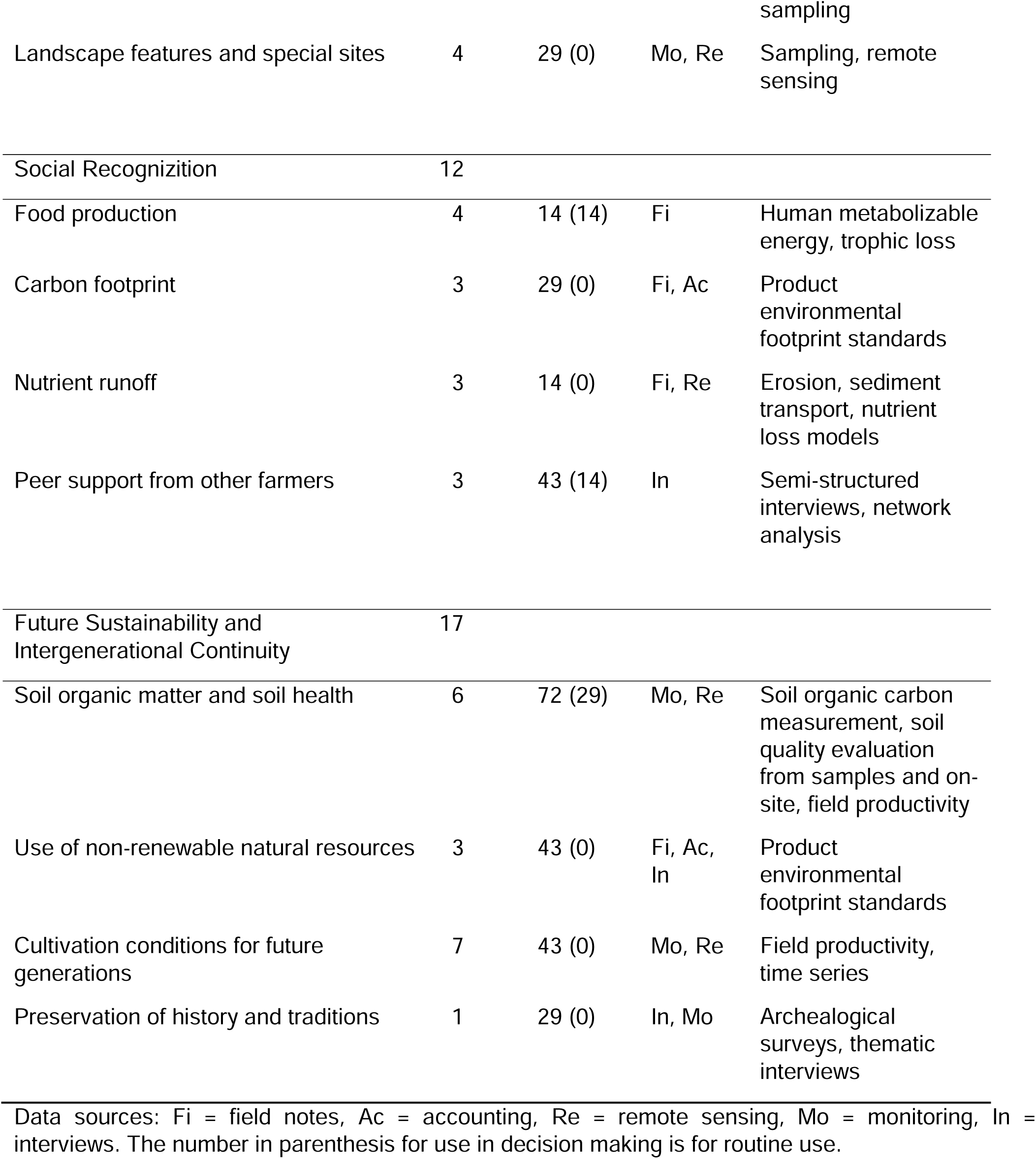
Decision making criteria, their average weight, use by interviewed farmers, and the workflow to calculate them.

Biodiversity criteria had relatively high priority weights but were not in routine use. This makes them a key target for further operationalisation. Currently they need on-site monitoring, which can become labour intensive when dealing with hundreds of fields. The same labour intensiveness problem also applied to future sustainability and next-generation cultivation conditions. Although soil health can be estimated from remote sensing, a more detailed study requires on-site monitoring, which is hard to scale to farm level. The social recognition subobjective group had the most developed methods for estimating carbon footprints or nutrient runoff, but also the lowest priority for on-farm decision making. Overall, there is a paucity of operational method development for on-farm decision making, especially for future generation cultivation conditions, biodiversity and organisation of work.

## 4. DISCUSSION

Farm-tailored and context-specific practices are essential for regenerative agriculture, but there is very little guidance on how to identify specific issues, and where to focus development on a farm. We have demonstrated that farmer values and objectives can be mapped with a simple questionnaire, highlighting individual preferences and common priorities. We also showed how open satellite monitoring data and GPP models can be used to find out how a farm compares to other farms in terms of productivity, and how the fields on the farm relate to each other. Importantly, we also used open remote sensing data to identify local erosion and soil loss hotspots, which comprise only 10% of the field area, but cause over half of the soil loss. These are critical steps in building an operational version of a farm context for decision making. Finally, we outlined how additional decision-making criteria could be operationalized using a combination of remote sensing, farm data, and monitoring.

Despite differences in scale and forms of production, the farms also had similarities in shared values. Our findings showed that farmers, contrary to stereotypes, value benevolence, universalism, and self-direction over power, achievement or tradition corroborating earlier findings (Graskemper et al., 2022; Sorvali et al., 2022). Self-transcendence and openness to change seemed to be strong values when compared with the general Finnish population (ESS, 2024) and in agreement with previous studies on farmer’s values (Graskemper et al., 2022; Sorvali et al., 2022). Like others we also found that farmers care about ecological matters, but prioritize the practical issues of profitability and operational efficiency (Beacham et al., 2023; Pannell et al., 2006). Improving profitability and soil functions at the same time is a key management target for regenerative agriculture, if it is to be interesting to farmers (Beacham et al., 2023; Jaworski et al., 2024; Schreefel et al., 2022).

Farmers may, however, face trade-offs between objectives. For example, the identified subset of high-productivity and high-erosion fields (section 3.4) presents a clear trade-off. To mitigate erosion, buffer strips and contouring might be necessary, but these would take very productive farmland out of production. Farmers’ goals and context determine whether they prefer to accept these trade-offs (Pannell et al., 2006). Farmers may be more willing to accept decrease in economic profit and production, if they place a strong weight on long-term soil improvement, decreasing erosion, or increasing biodiversity (Beacham et al., 2023). If the loss is considered too large, change in practices would require compensation from the government or supply-chain. To determine the required compensation level, several methods to determine farmer Willingness to Accept values have been developed (Boufous et al., 2023). One promising method is the reverse auction, where farmers submit financial bids to government agency indicating the compensation they require for implementing the practices (Ferraro, 2008; Latacz-Lohmann and Van der Hamsvoort, 1998). Since different fields and practices have different potentials for improving soil health or biodiversity, the most competitive bids are likely focused on field areas with the highest potential for improvement with the lowest impact on farming practices. The approach of this article demonstrates, how farmers could identify such priority areas and how government agencies could evaluate the bids, based on available open data and published models. Therefore, an operational context makes it easier for farmers to participate successfully in payments for ecosystem services schemes.

Contrary to the high-productivity field trade-offs with soil conservation, the low productivity and high variability fields present an opportunity for soil improvement. There was a 2–3-fold difference between the lowest and highest productivities. High and consistent productivity differences were also found within individual farms, indicating a great potential for increasing both productivity and carbon sequestration (Afshar et al., 2026; Mattila and Vihanto, 2024). The large interannual variation on variable fields indicated low resilience to extreme weather, which is a challenge for future climate mitigation (Afshar et al., 2026; Qiao et al., 2022). Diverse crop rotations and perennial crops were less sensitive to the interannual variation due to poor years.

The estimate of the 10% critical source area responsible for the majority of soil loss was in agreement with previous studies (Mattila and Niemi, 2026; McDowell et al., 2025). Erosion or sediment loss did not correlate with productivity, indicating management actions which improve productivity are not sufficient to manage soil loss (Mattila, 2024). Soil conservation measures should be targeted on subfield level, on local hotspots with high soil loss levels. The non-targeted soil conservation measures had already reduced erosion to approximately half of the worst-case scenario of annual crops and bare soil, but they were not sufficient on the high erosion zones. Farms with sloped fields and large catchments would need further conservation practices (contouring and buffer strips) as even the continuous plant cover promoted by regenerative agriculture cannot prevent excessive erosion on these sites.

Our demonstration had several limitations related to the small number of participating farmers, using only a single formulation of the decision criteria, and not quantifying biodiversity-related criteria. Our small sample size of 10 farms was sensitive to individual differences, but it nevertheless produced similar findings to larger surveys on soil management (n=478; Jaworski et al. 2024) or farmer values (n=787 and 4401; Sorvali et al. 2022; Graskemper et al. 2022). However, the low number of participants limited our possibilities for testing multiple versions of the decision-making criteria (Fig 1 and Table 1). As such the criteria list is one example of a multiple-objective farm development framework, which we believe could be refined and made more generalizable, by considering previous work on farmer value systems (Donovan et al., 2025; Hill and Bradley, 2024).

We were able to identify on-farm hotspots for erosion, but not for biodiversity. High resolution, spatial models have been developed for pollinators (Abdi et al., 2021; Gardner et al., 2020), but we were not able to parameterize them in the scope of this study, in particular to handle effects of in-field heterogeneity. The transferability of statistical models of pollinator densities in arable fields is limited (Blasi et al., 2021), and employing process-based pollinator models in a novel context without the availability of empirical datasets for model calibration may introduce substantial uncertainty (Gardner et al., 2020). Process-based models may nevertheless be very useful in the context of our study, and testing and contrasting these with models on productivity, erosion and profitability will be important future steps to assess the potential of regenerative farming methods to deliver multifunctional benefits (Winberg et al., 2026), and ultimately to develop tools for decision support on the farm level.

Overall, we were able to test and bring together value elicitation, field productivity assessment, and erosion mapping to present an outline of a decision-making context on the farm scale. Our assessment considers which fields underperform, which are climate resilient and productive, and how a specific farm relates to other similar farms. Although many of the criteria were left unquantified, we provide a first draft of a context that can guide where to focus further work and which parts of farm development to focus on.

## 5. CONCLUSIONS

Although context-specific and localized solutions are essential for the effective application of regenerative agriculture, defining the context has been challenging until now. Here we demonstrated that to operationalize the definition of a context, one can map values and decision-making criteria effectively with a weighting questionnaire. Fields and farms can be benchmarked based on their productivity to identify problem fields and room for improvement. Finally, high resolution digital elevation models enable identifying erosion and sediment hotspots, which should be addressed for long term soil health. Data and models are increasingly available to support on-farm decision making. Linking those through operational workflows is key for helping farmers manage and even reverse soil degradation on their farms.

## Supplementary information

**Figure S1.**
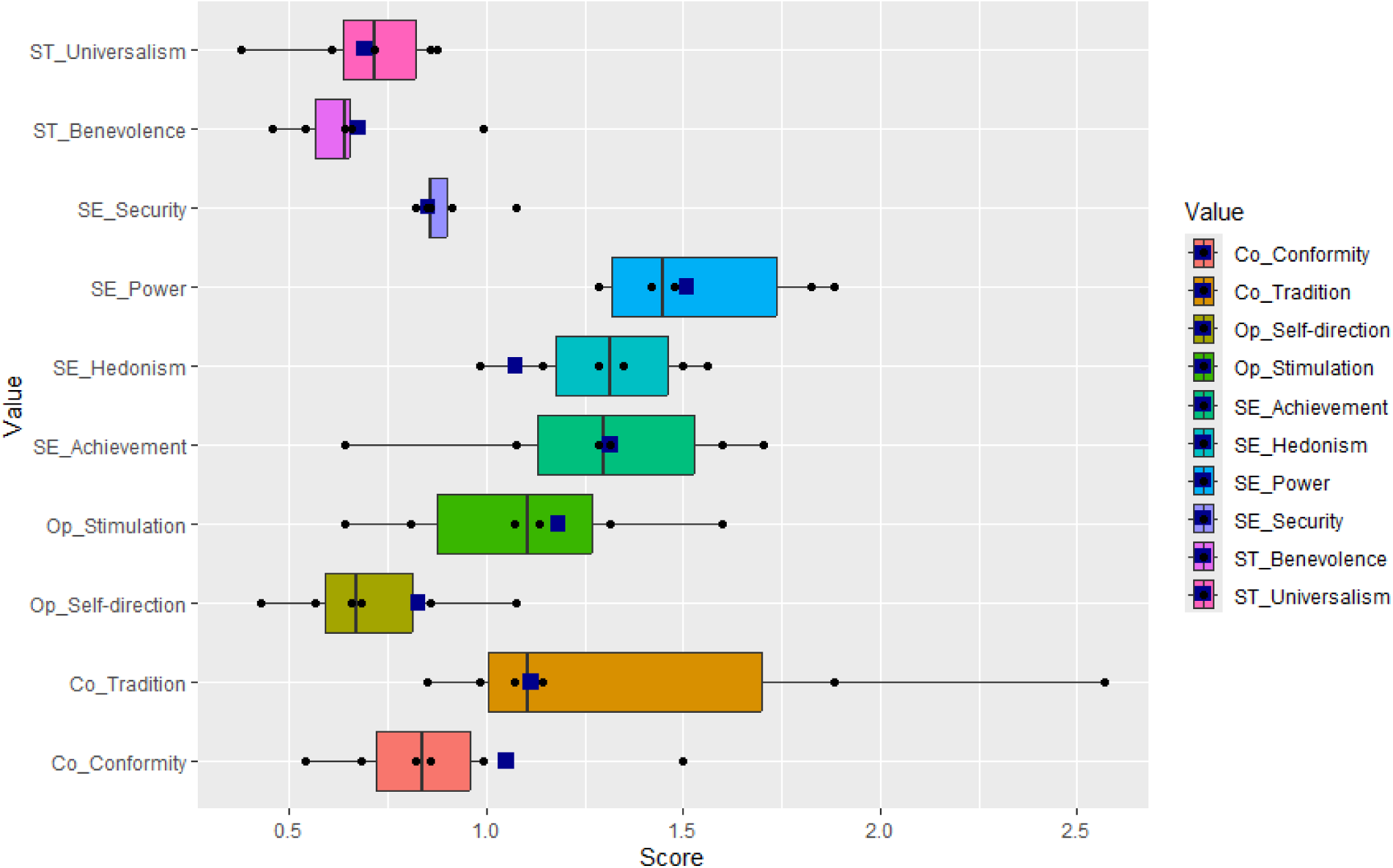

**Figure S2.**
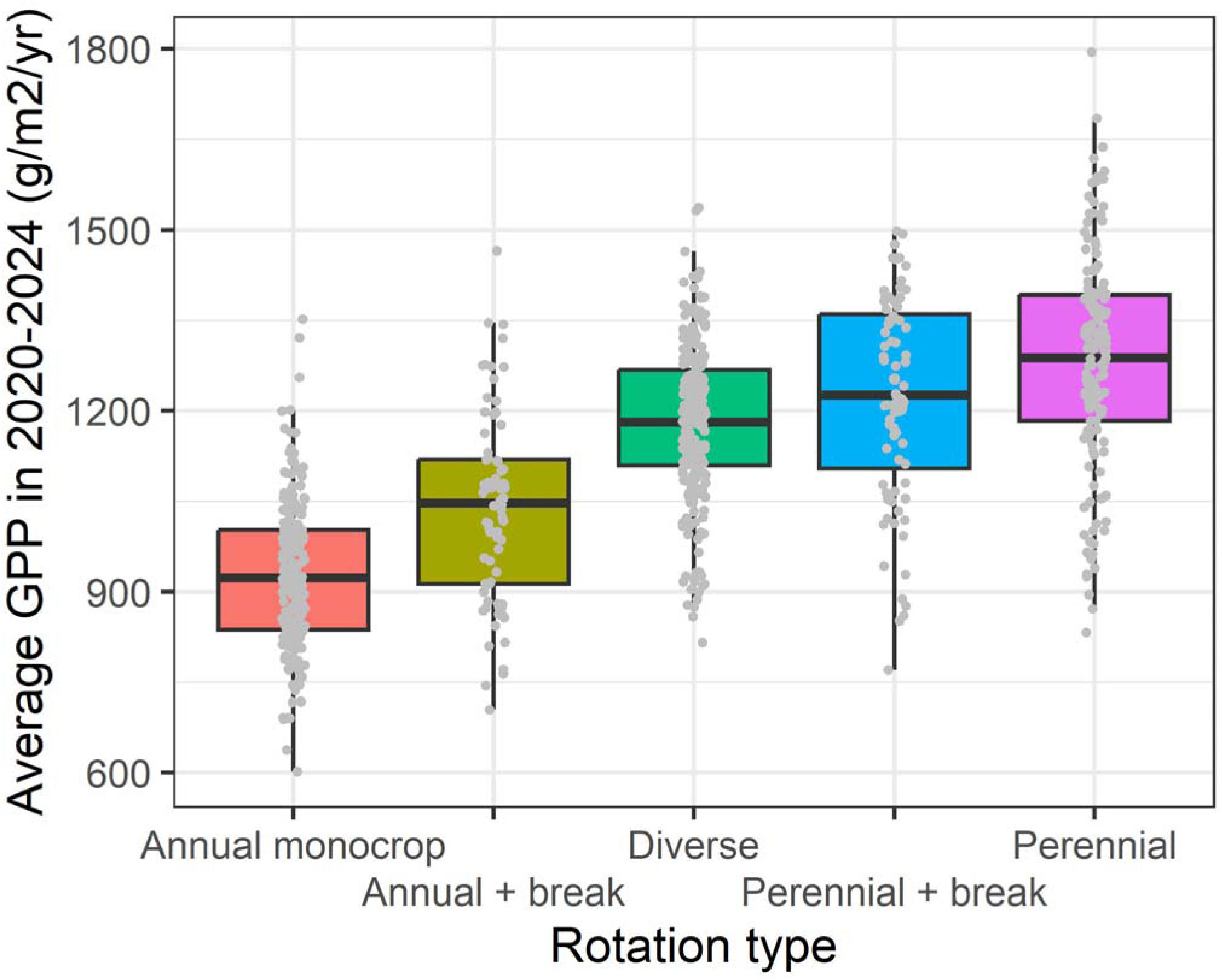

**Figure S3.**
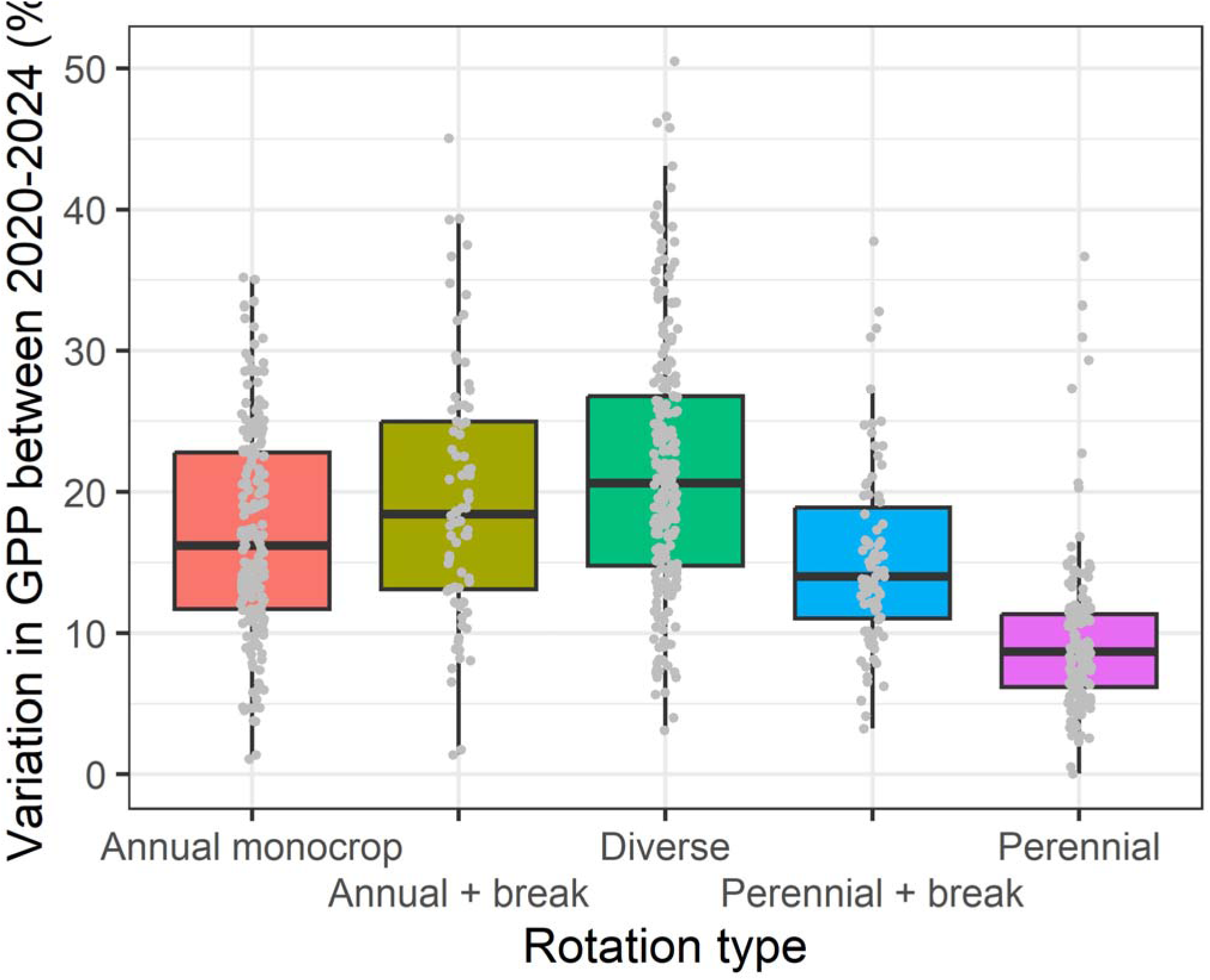

**Figure S4.**
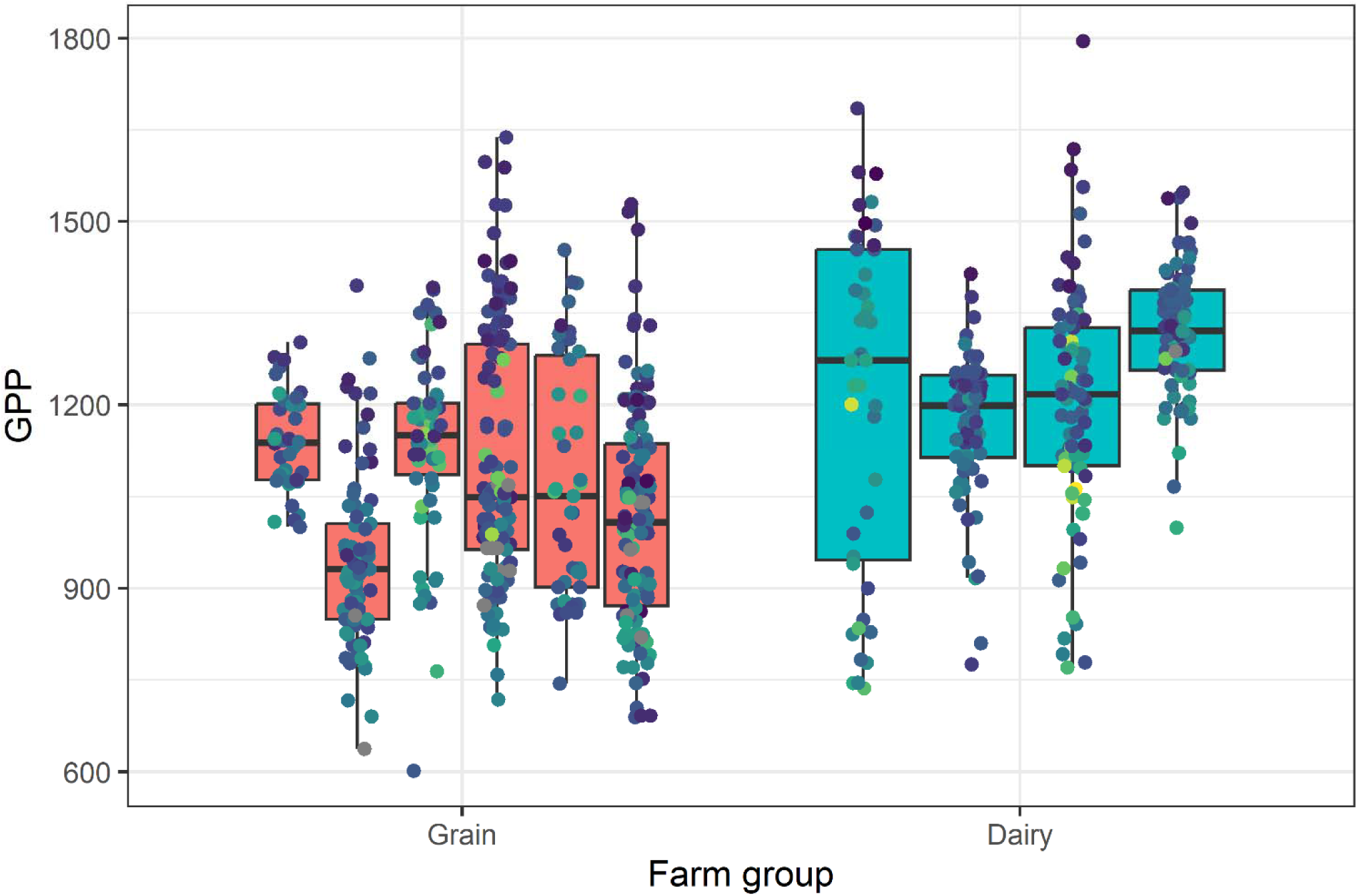

**Figure S5.**
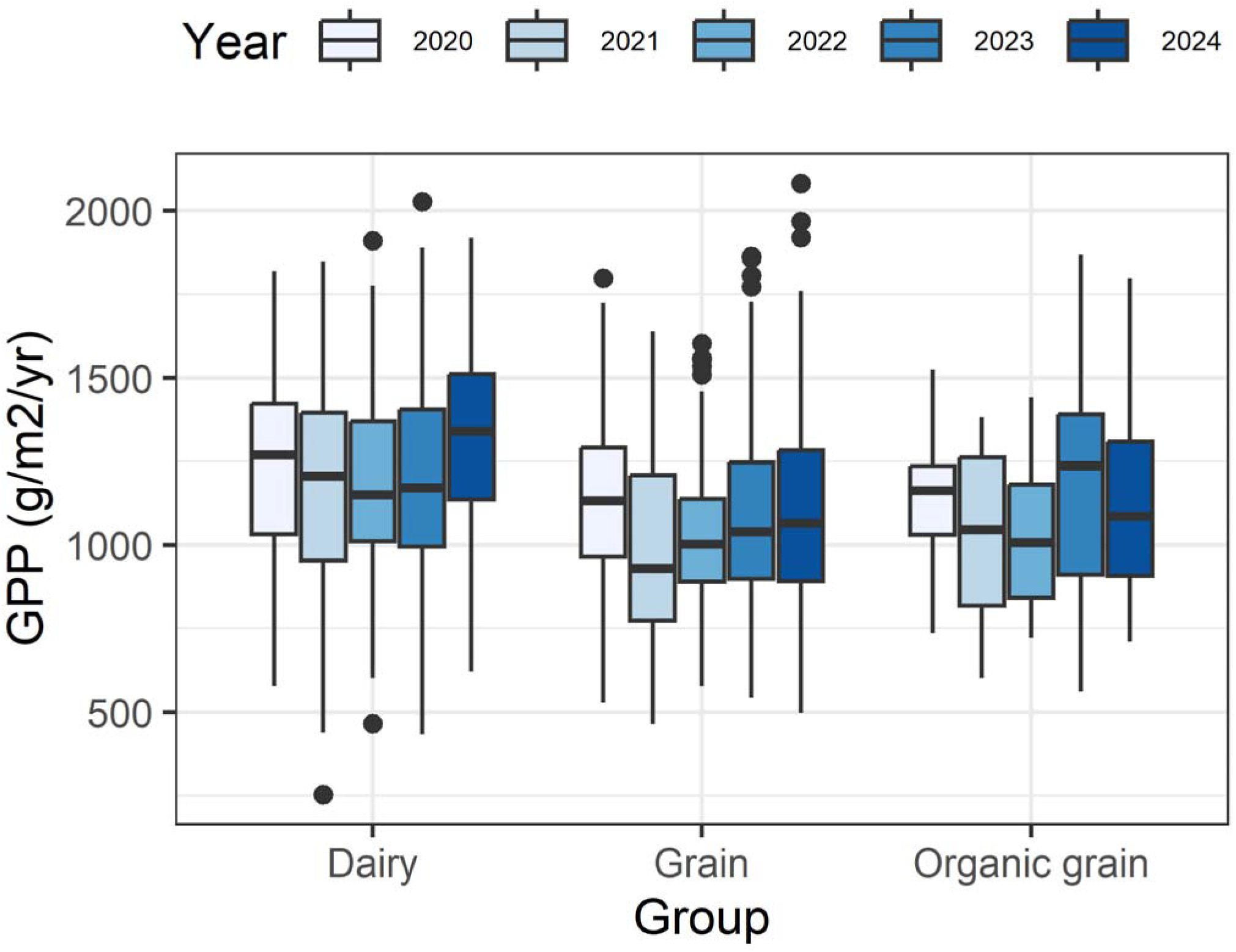

## Notes

### Competing Interest Statement

The authors have declared no competing interest.

### Funding

- Ministry of Agriculture and Forestry: va-mmm-2025-9-38

