## Supplementary figures for "Operationalising the context in regenerative agriculture: decision-making and farm variability"

Supplementary information

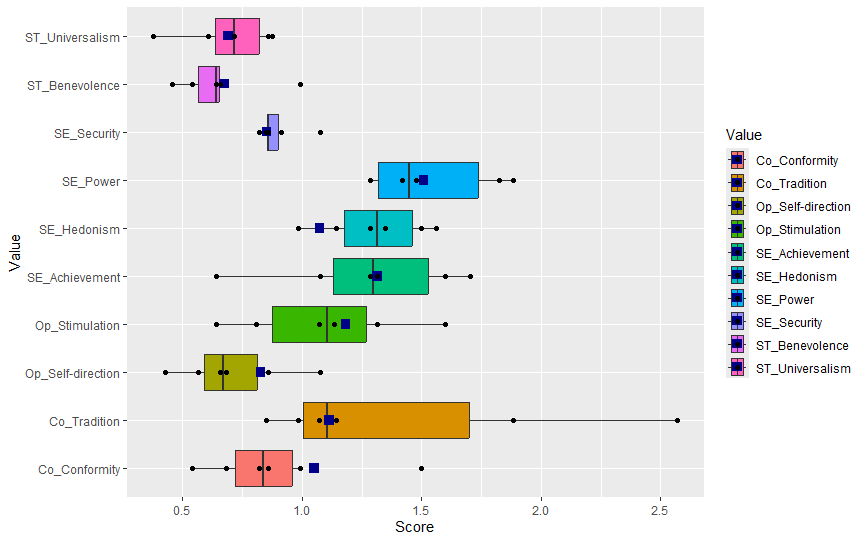

Figure S1. Farmer values as represented by the universal 10 values in the European Social survey. The blue square represents average Finnish values, while the boxplot and points are the responding farmer answers.

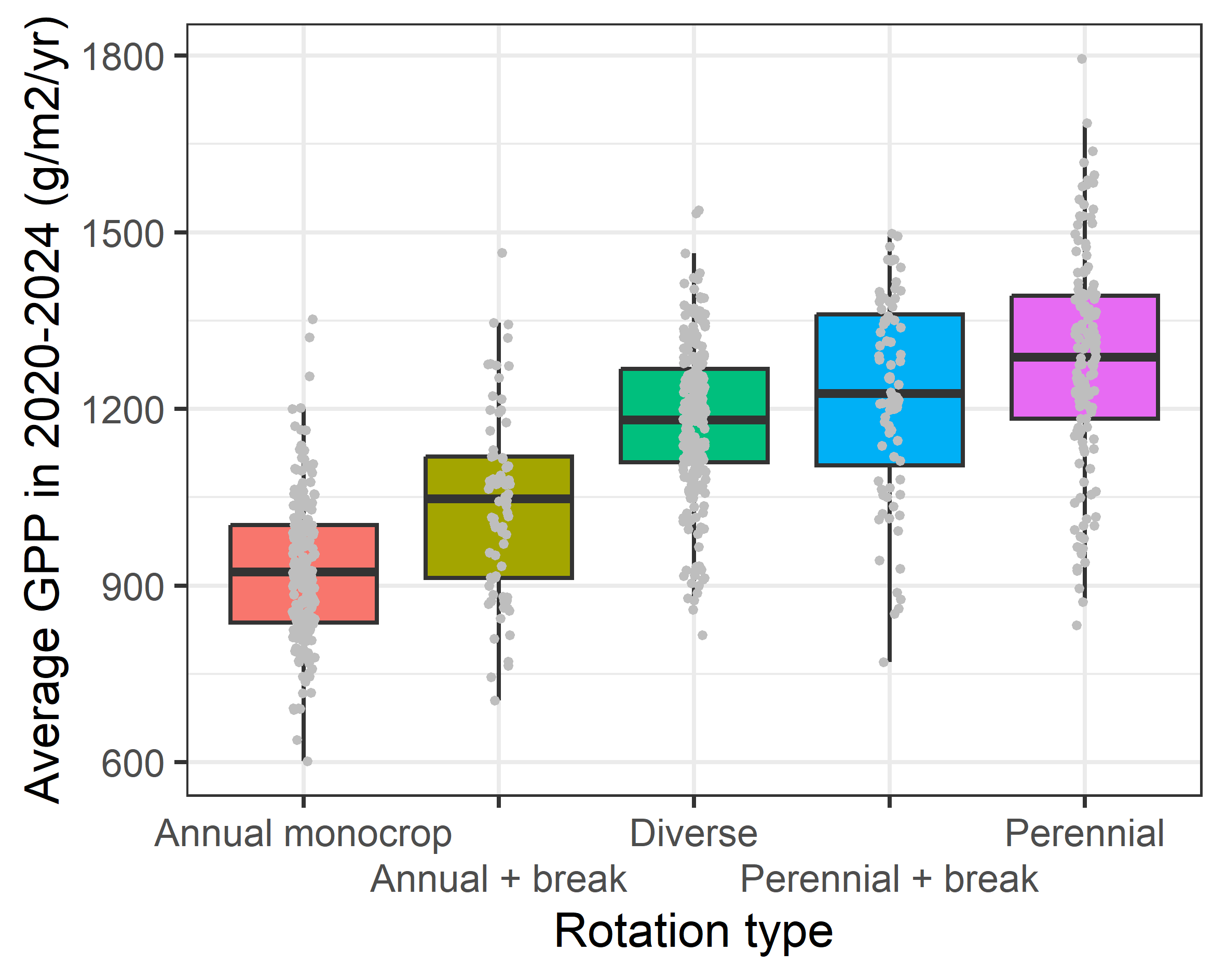

Figure S2. Field average productivity over five years classified by the crop rotation type on the field. Break implies one year of five, while diverse implies more than one year of five in either perennial or annual crops.

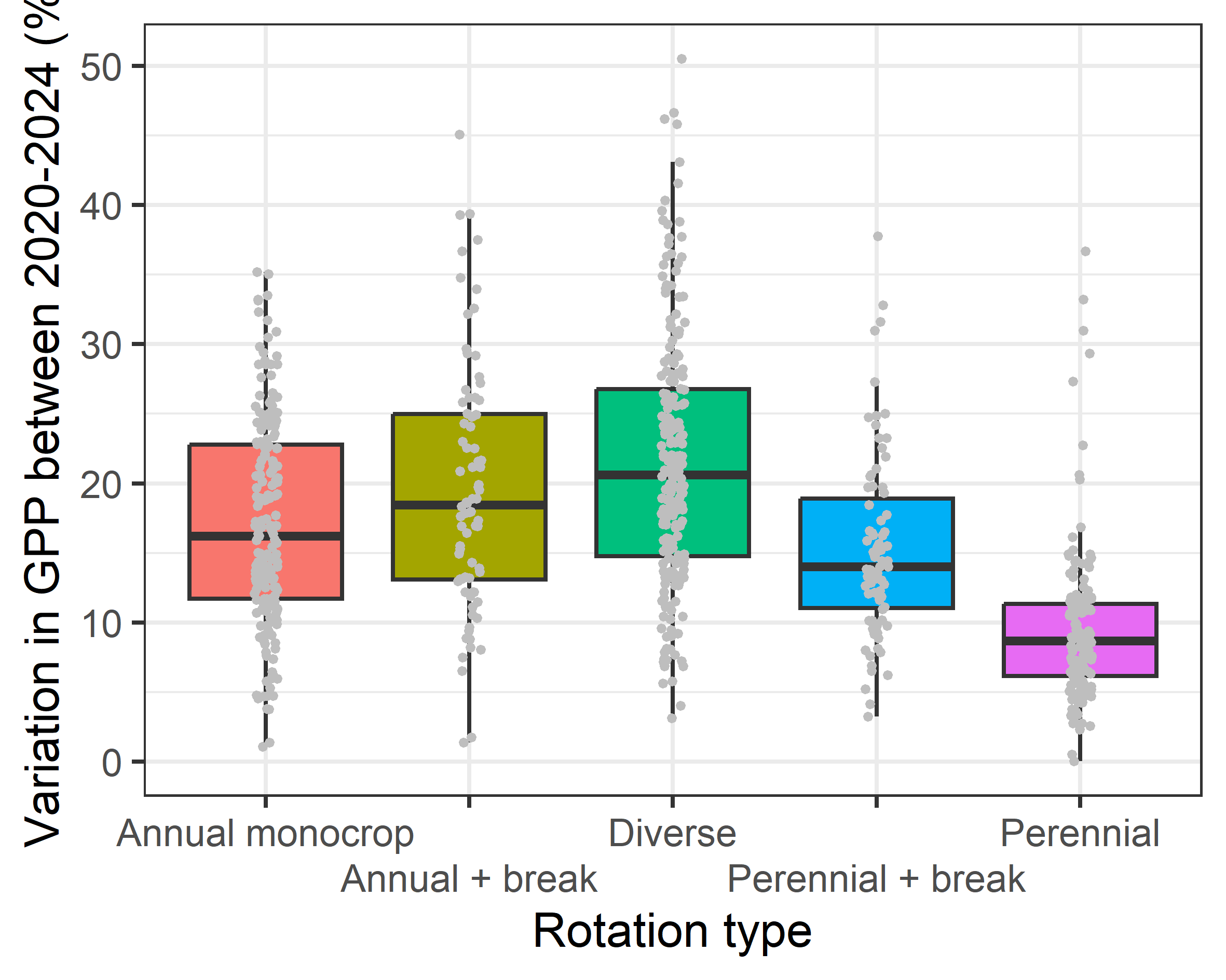

Figure S3. Variation in field productivity over five years classified by the crop rotation type on the field. Break implies one year of five, while diverse implies more than one year of five in either perennial or annual crops.

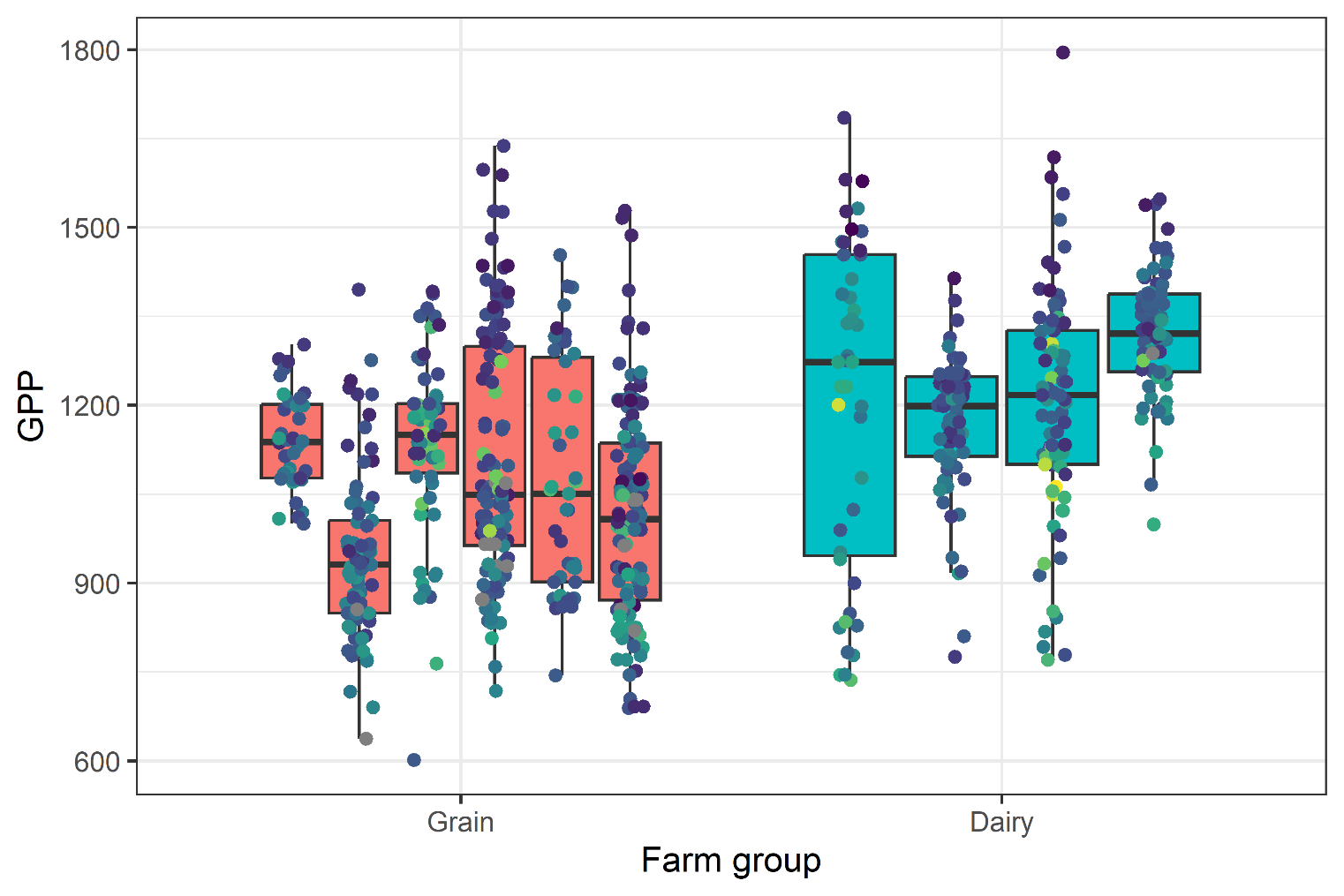

Figure S4. Average productivity (GPP g C/m2/yr) over five years per field as grouped by farms and farming systems.

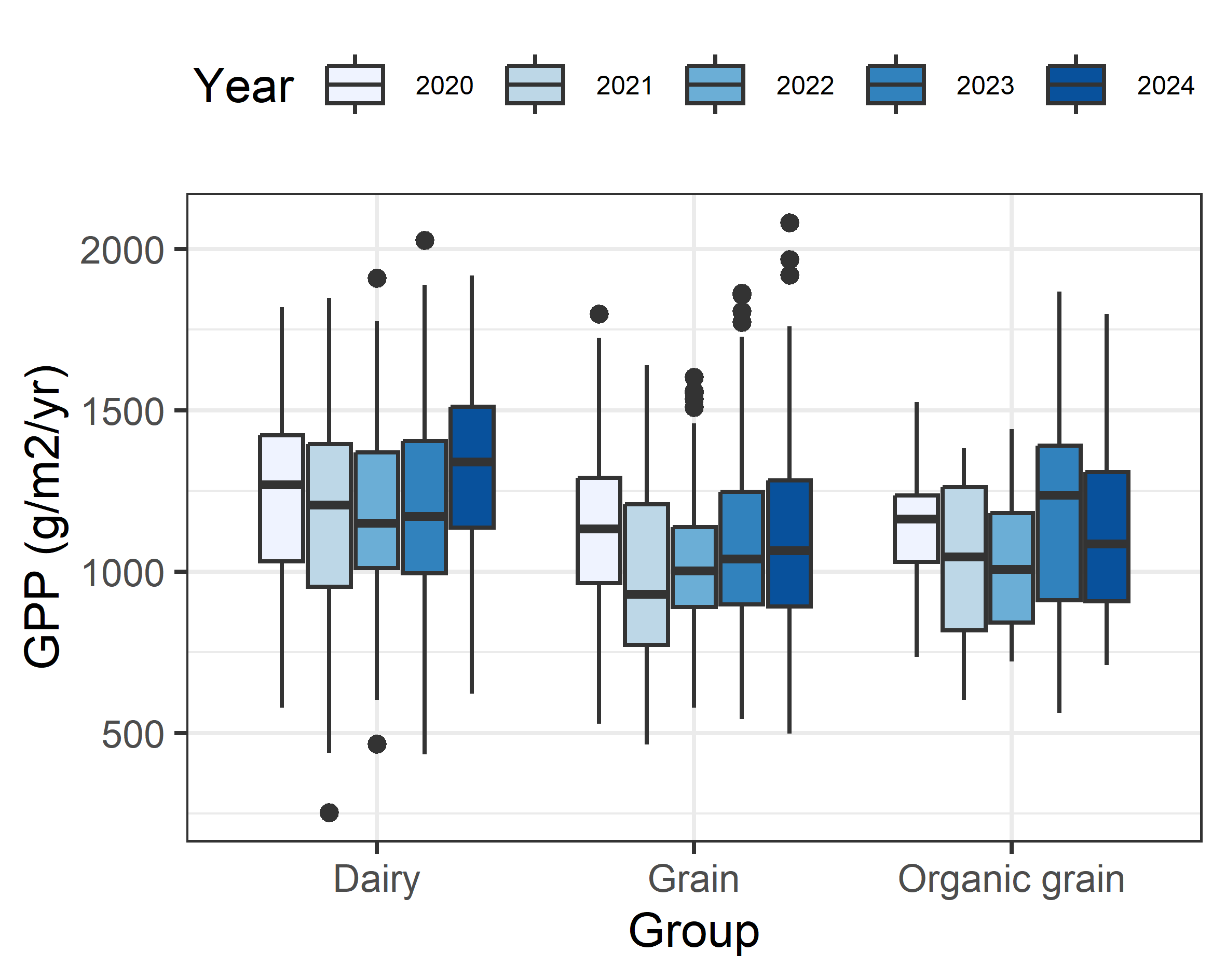

Figure S5. Annual productivity over five years in three different farm groups: dairy farms, conventional grain farms and organic grain farms. Years 2021 and 2022 were significantly weaker in grain farms but not in the two others.
